# Depression of heart rate in fish at critically high temperatures is due to atrioventricular block

**DOI:** 10.1101/2020.03.17.994947

**Authors:** Jaakko Haverinen, Matti Vornanen

**Affiliations:** University of Eastern Finland, Department of Environmental and Biological Sciences

**Keywords:** fish heart, electrocardiogram, cardiac arrhythmia, sodium current, chronaxie, rheobase

## Abstract

At critically high temperature, cardiac output in fish collapses due to depression of heart rate (bradycardia). However, the cause of bradycardia remains unresolved. Here we provide a mechanistic explanation for the temperature induced bradycardia. To this end rainbow trout (*Oncorhynchus mykiss*; acclimated at +12°C) were exposed to acute warming, while cardiac function was followed from electrocardiograms. From +12°C to +25.3°C, electrical excitation between different parts of the heart was coordinated but above +25.3°C atrial and ventricular beating rates became partly dissociated due to 2:1 atrioventricular (AV) block. With further warming atrial rate increased to the peak value of 188 ± 22 bpm at +27°C, while the rate of the ventricle reached the peak value of 124 ± 10 bpm at +25.3°C and thereafter dropped to 111 ± 15 bpm at +27°C. In single ventricular myocytes, warming from +12°C to +25°C attenuated electrical excitability as evidenced by increases in rheobase current and critical depolarization required to trigger action potential. The depression of excitability was caused by temperature induced decrease in input resistance (sarcolemmal K^+^ leak via the outward I_K1_ current) of resting myocytes and decrease in inward charge transfer by the Na^+^ current (I_Na_) of active myocytes. Collectively these findings show that at critically high temperatures AV block causes ventricular bradycardia which is an outcome from the increased excitation threshold of the ventricle due to changes in passive (resting ion leak) and active (inward charge movement) electrical properties of ventricular myocytes. The sequence of events from the level of ion channels to the cardiac function *in vivo* provides a mechanistic explanation for the depression of cardiac output in fish at critically high temperature.

## INTRODUCTION

Physical performance of fish is dependent on the oxygen supply to the tissues, which is largely determined by the quantity of blood delivered to them. The volume of blood circulated through the body in a unit of time is cardiac output which is the product of heart rate (*f*_H_), or the number of heart beats per minute, and stroke volume, or the volume of blood pumped by the ventricle in one contraction. In fish, acute warming initially increases cardiac output but at temperatures slightly below the critical thermal maximum of the animal cardiac output first plateaus and then sharply collapses (Stevens et al., 1972). The temperature induced collapse of cardiac output is caused by depression of *f*_H_ (bradycardia), since stroke volume is almost independent of temperature (Randall, 1968; Gollock et al., 2006; Sandblom and Axelsson, 2007; Steinhausen et al., 2008; Clark et al., 2008; Mendonca and Gamperl, 2010; Ekström et al., 2014; Penney et al., 2014; Ekström et al., 2016; Motyka et al., 2016; Ekström et al., 2019). Despite several studies, the mechanism of the temperature induced bradycardia remains unknown. This is a large gap in a fundamental physiological issue, because limitations in oxygen delivery to the tissues is considered to set the thermal tolerance limits of fish (Lannig et al., 2004; Farrell, 2009). Several speculations about the mechanistic basis of the temperature induced bradycardia have been suggested but thus far no definitive experimentally verified explanation have been provided. Since the mechanism of bradycardia remains unknown, there is also uncertainty about its physiological role: is it useful (adaptive) or harmful (non-adaptive) trait?

Rate and rhythm of the heart beat is determined in the sinoatrial pacemaker and therefore an obvious candidate for bradycardia is the direct temperature-dependent slowing of the primary pacemaker rate (Haverinen and Vornanen, 2007). On the other hand, temperature-induced bradycardia may in principle be analogous to hypoxic bradycardia, which improves oxygenation of the myocardium (Farrell, 2007). In the later scenario high temperature results in hypoxemia and activation of oxygen sensitive chemoreceptors which trigger a controlled (adaptive) bradycardia via the cholinergic system (Hughes and Roberts, 1970; Rantin et al., 1998; Gollock et al., 2006; Ekström et al., 2016), or alternatively the effect is mediated by thermal receptors (Hughes and Roberts, 1970). Also the possibility, that temperature induced elevation of external K^+^ concentration (hyperkalemia) suppresses the pacemaker rate, has been raised (Heath and Hughes, 1973). One more putative explanation is that acute rise of 4 temperature changes the fluidity of the lipid membrane and impairs Na^+^, Ca^2+^ and K^+^ ion passage through the sarcolemma (Lennard and Huddart, 1991).

The above alternatives, with the exception of the last one, suggest that the bradycardia is regulated at the level of the *sinus venosus* or the atrium. However, i*n vitro* force recordings from fish hearts for almost 90 years ago show that during acute warming the ventricle stops working first, then stops the atrium, whereas the sinoatrial pacemaker is best able to withstand high temperatures (Markowsky, 1933; Koehnlein, 1933; von Skramlik, 1935). However, it has not been directly tested whether the temperature-induced bradycardia is due to (i) the suppression of the supraventricular tissues (sinoatrial pacemaker, atrium) or (ii) failure of the ventricular excitation. This problem is possible to solve by locating the site of bradycardia among the several specialized compartments of the fish heart. If the former hypothesis (i) is true, then P wave (atrial depolarization) and QRS complex (ventricular depolarization) should be always coupled and atrial and ventricular beats should slow down in the same rhythm. Conversely, if the latter hypothesis (ii) is correct P wave and QRS complex should occur partly independently of one another and beating rate of the ventricle should be lower than that of the atrium. Therefore, the first aim of the present study was to test the two alternative hypotheses of the temperature induced bradycardia by *in vivo* ECG recordings in rainbow trout (*Oncorhynchus mykiss*). A careful examination of ECGs showed that P wave and QRS complex became dissociated at critically high temperatures thus supporting the latter hypothesis. Thereafter, the second aim of the study was to seek a mechanistic explanation for the dissociation of atrial and ventricular depolarizations. The previous findings from brown trout (*Salmo trutta*) and roach (*Rutilus rutilus*) ventricular myocytes have shown that temperature dependencies of the Na^+^ inward current (I_Na_) and the inward rectifier K^+^ current (I_K1_) – two antagonistic ion currents critical for initiation of cardiac AP (Varghese, 2016) – are widely different. I_Na_ is depressed at relatively low temperatures, while I_K1_ is heat resistant and increases almost linearly with rising temperature (Vornanen et al., 2014; Badr et al., 2017). If the failure of the fish ventricle is due to temperature induced mismatch between depolarizing I_Na_ and repolarizing I_K1_, then the inward charge required to trigger AP should exceed the charge provided by I_Na_. To test this hypothesis, we measured the temperature dependence of inward charge transfer by I_Na_ and temperature dependence of the charge demand of the quiescent ventricular myocytes.

## MATERIALS AND METHODS

### Animals

Rainbow trout, 5.6 ± 17.8 g in body mass (mean ± SEM, *n*=37), were bought from the local fish farm (Kontiolahti, Finland). In the animal facilities of the university, the fish were maintained in 500 L metal aquaria for a minimum of 3 weeks before the experiments. Water temperature was regulated at +12 ± 0.5°C (Computec Technologies, Joensuu, Finland) and full oxygen saturation was maintained by aeration with compressed air. Ground water was constantly flowing through the aquaria at the rate of 150-200 L per day. Trout were fed 5 times per week *ad libitum* with trout feed (EWOS, Turku, Finland). The experiments were authorized by the national animal experimental board in Finland (permission ESAVI/8877/2019).

### Recording of electrocardiogram (ECG)

ECG recordings were made using the previously described methods (Vornanen et al., 2014; Badr et al., 2016). Anaesthetized trout (neutralized tricaine methane sulfonate, 0.3 mg/L, Sigma, St Louis, MO, USA) were placed ventral side up on an operating table and the gills were irrigated with circulating tap water. Two thin stainless-steel electrodes (diameter 0.23 mm; A-M Systems, Carlsborg, WA, USA) were obliquely inserted from the ventral surface close to the pericardium. The electrode wires were fixed on the ventral sided of the fish by two sutures. The fish was placed into a cylindrical Perplex chamber (length 29 cm, diameter 9 cm, volume 1.84 L) with open ends. The chamber and the reference electrode were immersed in a large (250 L) temperature-regulated (+12 ± 0.5°C) stainless steel aquarium with constant aeration of the water. The fish were allowed to recover from the operation for 1-2 days. The recovery was considered complete, when a clear and steady *f*_H_ variability appeared in the ECG. Temperature-dependence of ECG was studied by a procedure where water temperature was warmed at the constant rate of 3°C/h (Computec Technologies, Joensuu, Finland) starting from the acclimation temperature of the fish (+12°C). The experiment was discontinued when ventricular beating rate turned to steady decline. Immediately after the experiment the fish was killed by a strong blow to the head and pithing. During the experiment ECG and water temperature were continuously recorded via bioamplifiers (ML 136, ADInstruments, Colorado Springs, CO, USA) and the digital recording system (PowerLab, ADInstruments) on the computer. In ECG, P wave, QRS complex and T wave correspond atrial depolarization, ventricular depolarization and ventricular repolarization, respectively. The following parameters were measured from these waveforms: PQ interval (the time taken by impulse conduction from the beginning atrial contraction to the beginning of ventricular contraction), QRS duration (the time taken by impulse transmission through the ventricle) and QT interval (the average duration of ventricular AP) (Vornanen et al., 2014; Badr et al., 2016). *f*_H_ was separately calculated for the ventricle (*f*_HV_) and the atrium (*f*_HA_) from the number of QRS complexes and P waves, respectively.

### Patch-clamp methods

Ventricular myocytes were enzymatically isolated with the established method for fish hearts (Vornanen, 1997; Vornanen et al., 2002). Cell isolation was conducted at room temperature (21-22°C) and the isolated myocytes were stored at +5°C. Fresh cells were prepared in each experimental day.

The whole-cell current-clamp and voltage-clamp measurements were made using the standard methods and equipment of the whole-cell patch-clamp as reported previously in detail (Vornanen, 1997; Badr et al., 2018; Abramochkin et al., 2019). For recording of APs or ion currents, a small aliquot of cell suspension was placed into a chamber with a continuous temperature-controlled fluid flow through. After establishing a giga ohm seal and getting electrical access to the cell, transients due to series resistance (4-8 MΩ) and pipette capacitance were cancelled, and the capacitive size of ventricular myocytes was determined. APs and I_Na_ tracings were digitized at 10 kHz and low pass filtered at 5 kHz. Off-line analysis of the recordings was done using the Clampfit 10.4 (Molecular Devices, Saratoga, CA, USA) software package. The external solution used for AP recordings contained (mmol l^-1^) 150 NaCl, 3 KCl, 1.2 MgCl_2_, 1.8 CaCl_2_, 10 HEPES, 10 glucose, and pH adjusted with NaOH to 7.7 at 20°C. The composition of the internal saline solution was as follows (mmol l^-1^): 140 KCl, 5 Na_2_ATP, 1 MgCl_2_, 0.03 Tris-GTP, 10 HEPES (pH adjusted with KOH to 7.2 at 20°C) (Badr et al., 2018; Abramochkin et al., 2019). The same solutions were used for recording of the inward rectifier K^+^ current, I_K1_.The external saline solution for I_Na_ recordings composed of (mmol l^-1^) 20 NaCl, 120 CsCl, 1 MgCl_2_, 0.5 CaCl_2_, 10 glucose and 10 HEPES with pH adjusted to 7.7 with CsOH at 20°C. Nifedipine (10 μmol l^-1^, Sigma) was included to block I_Ca_. The pipette solution contained (in mmol l^−1^) 5 NaCl, 130 CsCl, 1 MgCl_2_, 5 EGTA, 5 Mg_2_ATP and 5 HEPES (pH adjusted to 7.2 with csOH at 20°C) (Haverinen and Vornanen, 2006).

The following AP parameters were analysed off-line: resting membrane potential (V_rest_, mV), threshold potential for AP initiation (V_th_, mV), threshold current (I_th_, pA), critical depolarization (cD = V_th_-V_rest_, mV), AP overshoot (mV), AP amplitude (mV), AP duration at 50% repolarization level (APD_50_, ms), maximum rate of AP upstroke (+dV/dt, mV ms^-1^) and the maximum rate of AP repolarization (-dV/dt, mV ms^-1^).

The following I_Na_ parameters were determined. The size of I_Na_ is expressed either as current density (pA pF^-1^) or charge transfer (pA ms^-1^ pF^-1^). The rate of I_Na_ inactivation is expressed as the time constant (τ) of the monoexponential fit (I_Na_ = a*(1-exp(−τ*t))) to the recorded current transients, where a is the peak amplitude of I_Na_, τ is time constant of inactivation and τ is time. Voltage-dependence of steady-state activation and inactivation were constructed by Boltzmann fits (V_SS-Act/SS-Inact_ = l/(1 + exp [(V - V_0.5_)/k]) to the experimental data. In the equation V_0.5_ is the half-voltage and the slope factor (k) for the voltage dependence. Recovery from inactivation was measured by the double pulse protocol, where the time distance between the pulses (from −120 mV to −20 mV for 50 ms) was stepwise increased.

### Input resistance and strength-duration relationship

Input resistance (R_in_) of isolated ventricular myocytes was measured at +12°C and +25°C under the current clamp mode of the patch-clamp using K^+^-based external and internal solutions (see *Patch-clamp methods*). Small depolarizing current pulses were injected into the cell at the V_rest_ of the cell and the changes of the membrane voltage were recorded. The time course of the voltage change was fitted using a mono exponential equation (see above) to obtain the time constant of the membrane, which is the product of membrane resistance and membrane capacitance (*τ* = R_in_ · C_m_). Knowing *τ* (ms) and the cell size (pF) R_in_ (MΩ) could be calculated from the equation.

The classical strength-duration relationship was determined under the same conditions as R_in_. The current (I) threshold for the initiation of AP was determined for square wave pulses of different durations (d, ms), which were delivered into the cell in random order at the rate of 1 Hz. The response of the cell was presented as a Weiss plot: the impulse strength is the charge (Q = I ·d, pA · ms^-1^) of the pulse (y-axis) which was plotted as a function of the pulse duration (x-axis) (Weiss, 1901; Geddes, 2004). This scatter plot was fitted by an equation for the straightline (Q= k + b*d), where k is the intersection point of the fitted line and the y-axis (the stimulus charge for infinitely short pulse) and b the slope of the line. The slope of the line (b) represents the rheobase, the smallest stimulus current that can induce AP when the pulse duration is infinitely long. The intersection of the fitted straight line and x-axis gives the chronaxie, the stimulus pulse duration required for initiation of AP when the current amplitude is equal to double the rheobase current (Irnich, 1980; Geddes and Bourland, 1985).

### Statistics

The results are given as means and the data variation is shown either as ± SEM or 95% confidence limits. Statistical differences between means were tested using t-test or Mann-Whitney *U* test. Tests were performed using IBM SPSS software (version 25.0). A *P*-value < 0.05 was considered to indicate a statistically significant difference between means.

## RESULTS

### Electrocardiogram (ECG)

EcG was recorded from conscious and slightly restrained fish with intact autonomic nervous regulation. QRS complexes and T waves were well reproduced in the ECG of all fishes (n=8), while P waves were clearly seen in the ECG of 4 fishes. At the acclimation temperature of the fish (+12°C) atrial and ventricular excitation were completely coordinated i.e. each P wave was always associated with QRS complex. However, at temperatures higher than +25°C atrial and ventricular excitations became partly dissociated in all 4 fishes where the P wave was clearly visible (Fig. 1A). Typically, every other P wave occurred without the associated QRS complex indicating a temperature-induced 2:1 atrioventricular (AV) block. With further warming this ended up in 3:1 or higher-level AV block. Because of the AV block, ventricular rate (*f*_HV_) increased only to the point where atrial and ventricular beating remained coordinated and then turned into bradycardia. The breakpoint temperature (*T*_BP_) for *f*_HV_ was +25.3 ± 0.4°C (Fig. 1A, B) and the peak value was 124 ± 10 bpm. Atrial beating rate (*f*_HA_), calculated from the number of P waves, steeply increased above the ventricular *T*_BP_ (Fig. 1A, B). The peak *f*_HA_ was significantly higher (188 ± 33 bpm at 27.0°C) than the peak *f*_HV_ (p<0.05). PQ interval (time taken by AP conduction through atrium and atrioventricular canal) and QT interval (mean duration of ventricular AP) became progressively shorter with warming. The duration of QRS complex (time taken by depolarization to spread through the ventricle) slightly increased at temperatures higher than +25°C (Fig. 1 C-E) (p>0.05).

**Figure 1.**
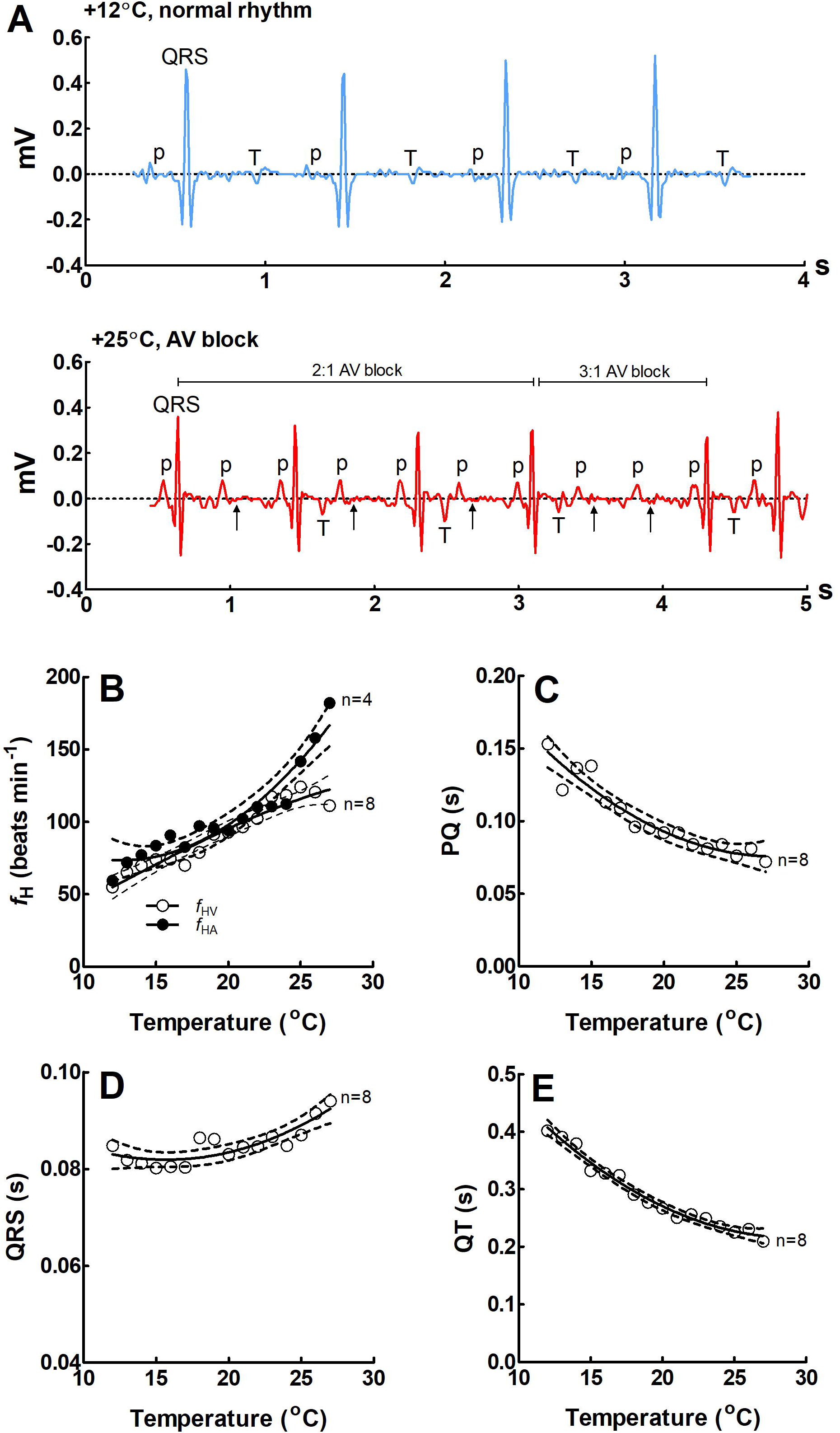
Effects of acute warming on ECG of rainbow trout. (A) Representative ECG recordings at +12°C and +25°C. The lower panel shows 2:1 atrioventricular block which ends up in 3:1 AV block (successive missing of two QRS complexes, small arrows). (B) Effect of temperature on atrial (*f*_HA_; n=4) and ventricular (*f*_HV_; n=8) beating rate determined from the number of P waves and QRS complexes, respectively. (C-E) Effect of temperature on PQ interval (C), QRS duration (D) and QT interval (E). Results are means of 8 fish. Dotted lines show ±95% confidence limits.

### Action potentials (AP)

To understand the cellular effects of high temperature, we measured the responses of ventricular AP to acute warming. APs were elicited by 4 ms current pulses of increasing strength at the rate of 1 Hz at +12°C and +25°C (Fig. 2A, B). Statistically significant differences (p<0.05) were found in 6 out of 9 AP parameters (Fig. 2c-F). V_rest_ and V_th_ became more negative at +25°C relative to +12°C but the change was bigger for V_rest_. Consequently, CD and I_th_ increased at +25°C. These findings indicate reduced EE of ventricular myocytes at +25°C. In addition, APD_50_ was strongly shortened and the amplitude of AP slightly enhanced at +25°C.

**Figure 2.**
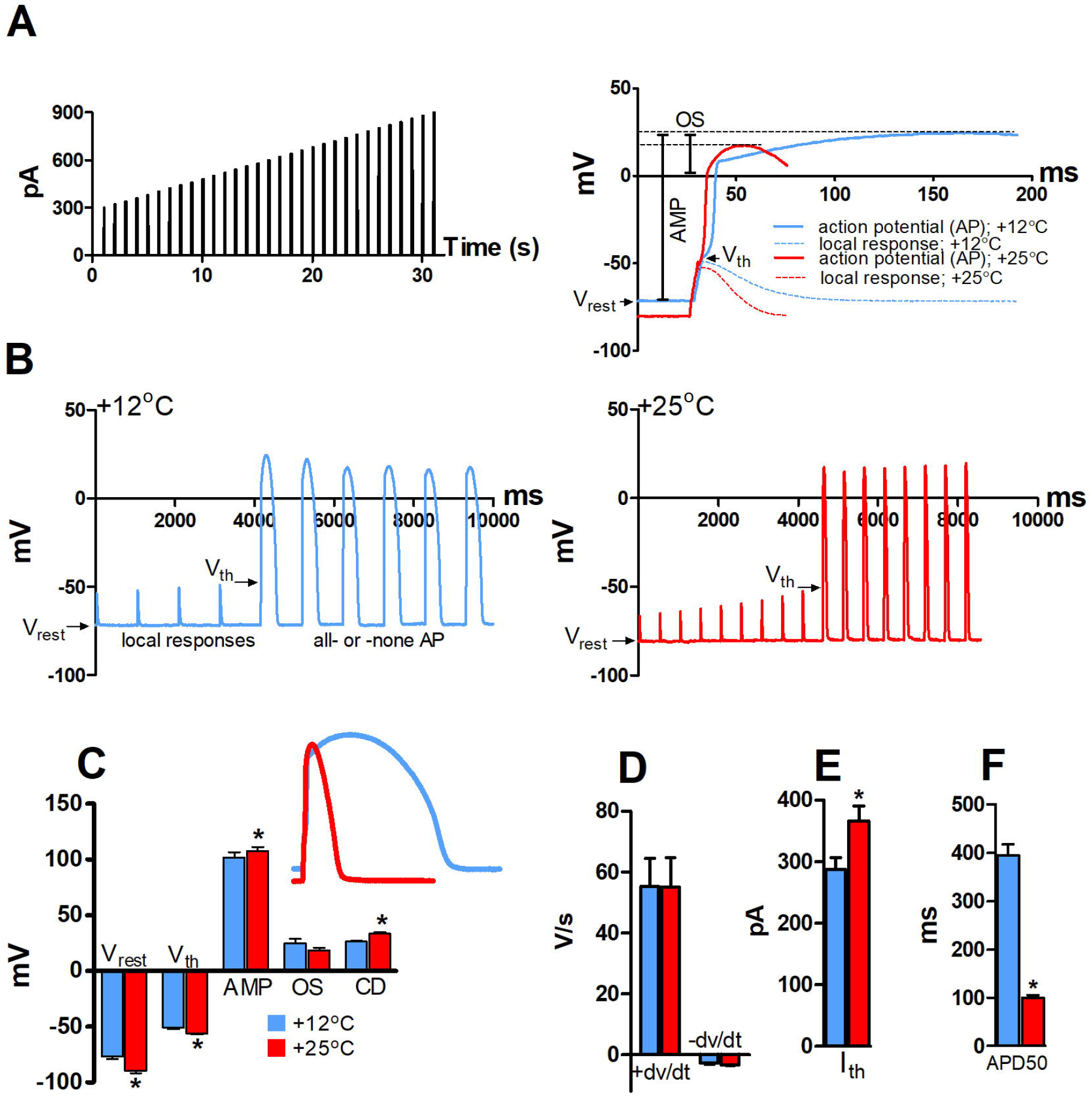
Effect of temperature on AP initiation of rainbow trout ventricular myocytes at +12°C and +25°C. (A) Stimulus current protocol (left) and representative fast sweep recordings of APs at +12°C and +25°C. The measured AP parameters are shown for the AP at +12°C (right). (B) Slow sweep recordings of ventricular APs +12°C (left) and +25°C (right) showing smaller voltage responses to stimulus current and a more negative V_th_ at +25°C compared at +12°C (left). (C) Effects of temperature resting membrane potential (V_rest_), threshold potential (V_th_), AP amplitude (AMP), AP overshoot (OS) and critical depolarization (CD). (D) Effects of temperature on the maximum rates of AP depolarization (+ dV/dt) and repolarization (-dV/dt). Effects of temperature on threshold current (I_th_) (E) and AP duration at 50% of repolarization (APD_50_) (F).

### Input resistance (R_in_) and strength-duration relationship

These experiments were done to reveal the charge requirement of AP initiation and its dependence on sarcolemmal ion leakage. R_in_ is a measure for the ion leak of the plasma membrane of quiescent ventricular myocytes and it was measured from the same cells at +12°C and +25°C. At +12°C the R_in_ was 2.1 times larger (528 ± 59 MΩ) than at +25°C (253 ± 28 MΩ) (p<0.01) indicating a leakier sarcolemma of ventricular myocytes at +25°C (Fig. 3A).

**Figure 3.**
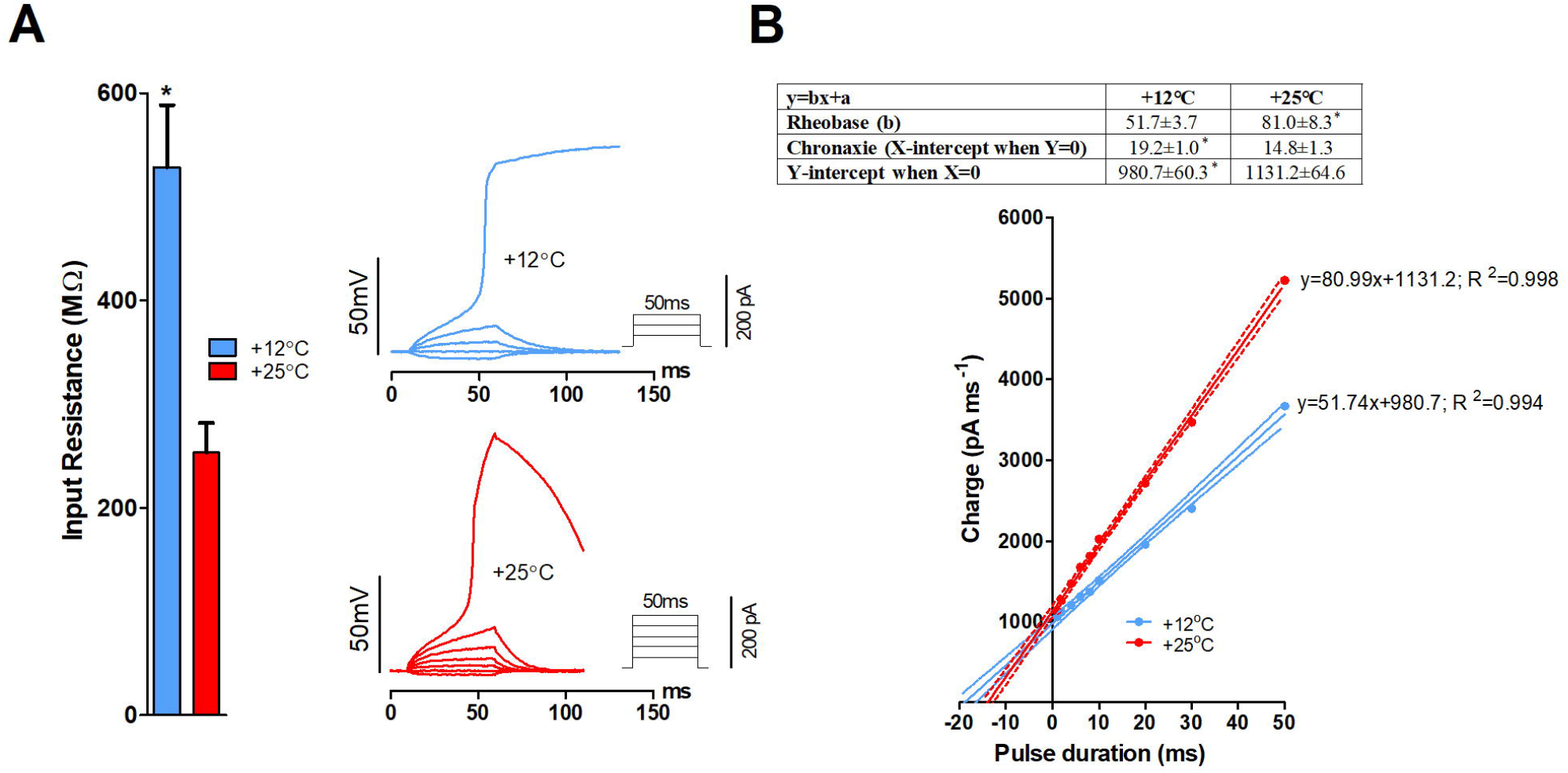
Effect of temperature on input resistance (R_in_) and strength-duration relationship of rainbow trout ventricular myocytes. (A) R_in_ of resting ventricular myocytes at +12°C and +25°C. (B) Effect of temperature on strength-duration relationship of ventricular myocytes. Representative current-clamp recordings showing the requirement of increased stimulus current/charge for AP initiation of the same myocyte at +25°C relative to that at +12°C (left). Mean results of strength-duration relationship at +12°C and +25°C presented in the form of Weiss plots (right) (n=13 myocytes from 4 fishes). The dotted lines indicate ± 95% confidence limits. The mean values (± SEM, n=13) for rheobase, chronaxie and the charge for infinitely short pulse (y intercepts of the lines) are shown in the box above the plot.

The inward charge transfer needed to bring V_rest_ to the V_th_ was examined at +12°C and +25°C by determining the classical stimulus strength-duration relationship in ventricular myocytes. Amplitude and duration of the current pulse were varied in random order and the charge needed to trigger AP at different pulse durations was plotted as a function of the pulse duration (Weiss plot) (Fig. 3B). At all pulse durations the charge needed to trigger AP was higher at +25°C than at +12°C (p<0.05). The rheobase current was 36.2% higher at +25°C (81.0 ± 8.3 pA) than at +12°C (51.7 ± 3.7 pA) (p<0.05), whereas the chronaxie duration was significantly shorter at +25°C (14.8 ± 1.3 ms) than at +12°C (19.2 ± 1.0 ms) (p<0.05). These data show that a larger absolute threshold current/charge is needed to trigger AP at +25°C than at +12°C, while shorter pulses are more effective at +25°C.

### Sodium current (I_Na_) and inward rectifier K^+^ current (I_K1_)

In these experiments the charge provided by the active ventricular myocytes and the resting leak via I_K1_ were measured. Current-voltage relationship of I_Na_ was determined at +12°C and +25°C. The peak density of I_Na_ was 20% larger at +25°C (−70 ± 4 pA pF^-1^) than at +12°C (−56 ± 3 pA pF^-1^) (p<0.05) (Fig. 4A, B). In contrast, the charge transfer i.e. the time integral of I_Na_ was 31% smaller (−40 ±2 pA ms^-1^ pF^-1^) at +25°C than at +12°C (−58 ± 3 pA ms^-1^ pF^-1^) (p<0.001) (Fig. 4C). *T*_BP_s of I_Na_ and I_K1_ were sought by gradual increases of the experimental temperature (3°C min^-1^) while the changes in current amplitudes/charge transfer were recorded (Fig. 4D, F). *T*_BP_s were +18.3 ± 0.6°C and +34.0 ± 1.1°C for the density of inward I_Na_ and outward I_K1_, respectively (p<0.001). Notably the charge transfer by I_Na_ started to decline immediately upon warming (Fig. 4E). The density of I_Na_ had declined to its starting level at +12°C when temperature was +27°C. At the *T*_BP_ of *f*_HV_ the charge transfer by I_Na_ was declined by 46%. Collectively this shows that supply of the charge decreases, and demand of the charge increases at high temperature.

**Figure 4.**
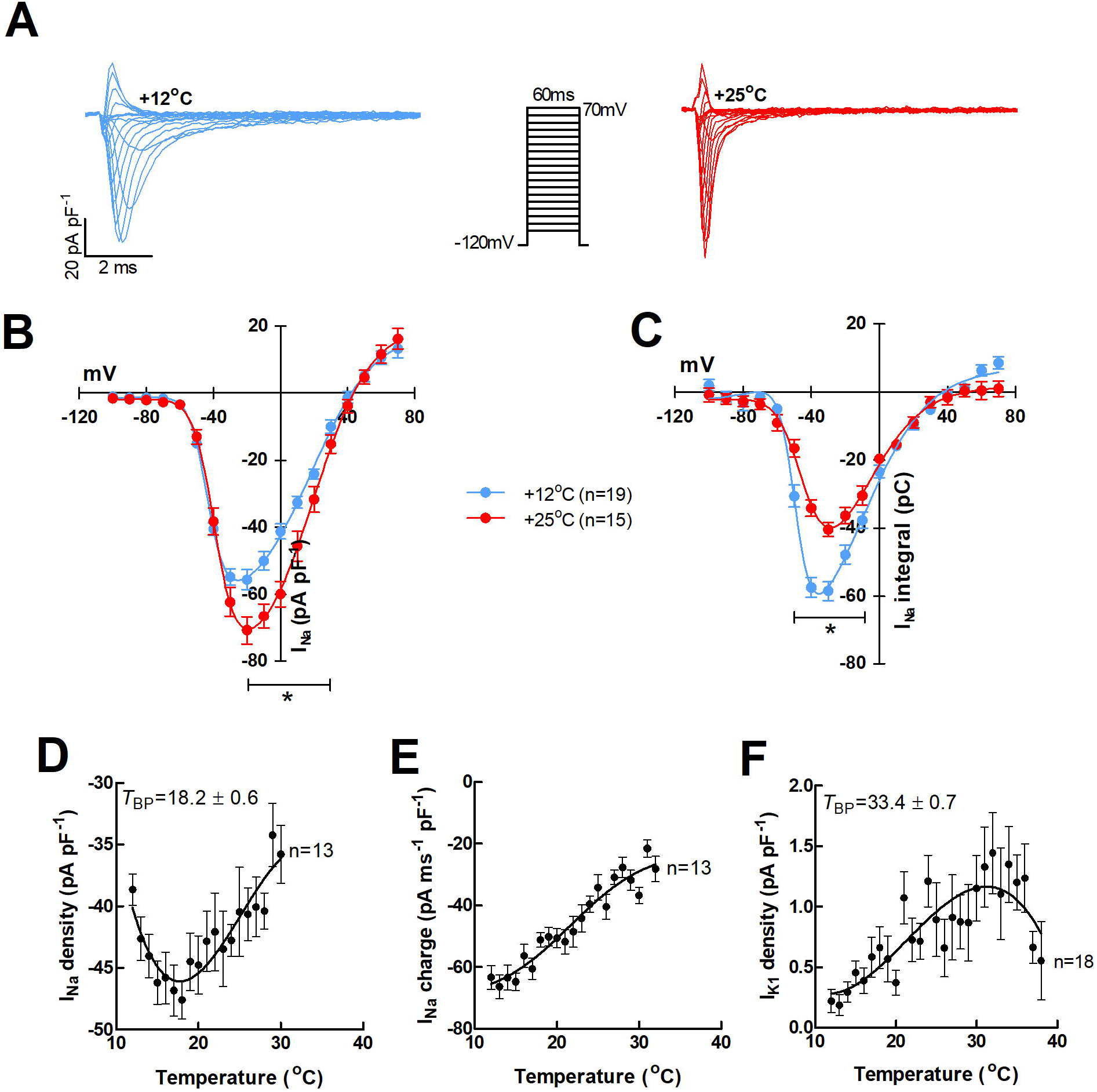
Effects of temperature on I_Na_ and I_K1_ of rainbow trout ventricular myocytes. (A) representative tracings of the voltage-dependence of I_Na_ at +12°C and +25°C. (B) Voltagedependence of I_Na_ density at +12°C (n=19 cells from 5 fish) and +25°C (n=15 cells from 5 fish). (c) Voltage-dependence of I_Na_ charge transfer at +12°C (n=19 cells from 5 fish) and +25°C (n=15 cells from 5 fish). (D and E) Effects of acute warming on I_Na_ density (D) and charge transfer (E). The results are means ± SEM of 13 cells from 3 fishes. (F) Effect of acute warming on the density of the outward I_K1_ in trout ventricular myocytes (n= 18 cells from 4 fishes).

Effect of warming on biophysical properties of I_Na_ was examined at +12°C and +25°C. Acute warming did not have effect on the voltage-dependence of steady-state inactivation but shifted steady-state activation curve by 4 mV to more positive voltages (p<0.05) (Fig. 5A). The rate of current inactivation became faster at +25 °C (Fig. 5B). Surprisingly, the rate of I_Na_ recovery from inactivation was not affected by acute temperature change (p>0.05) (Fig. 5C).

**Figure 5.**
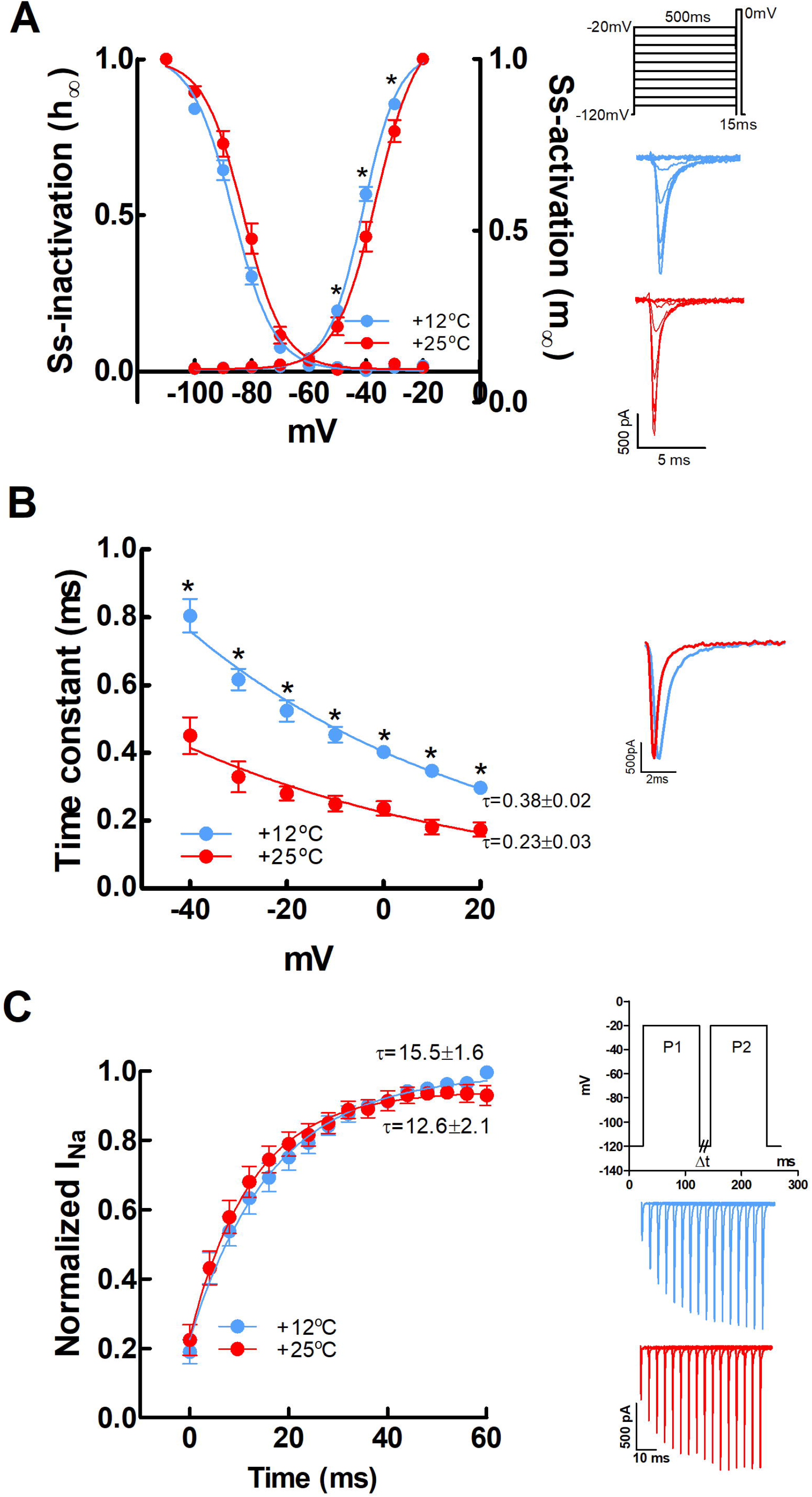
Temperature-dependence of I_Na_ in rainbow trout ventricular myocytes. (A) Voltagedependence of steady-state activation shifted more positive voltages (p<0.05) but steady-state inactivation was not affected (p>0.05) by acute increase in temperature from +12°C to +25°C. Mean results ± SEM of 19 cells from 5 animals (left). Representative recordings of I_Na_ at +12°C and +25°C. (B) Acute increase in temperature accelerated the rate of I_Na_ inactivation. Mean values ± SEM for the voltage-dependence of I_Na_ inactivation (left) and representative I_Na_ recordings at −20 mV) (right) at +12°C and +25°C. (C) Recovery of I_Na_ from inactivation. Acute increase of temperature from +12°C to +25°C did not have any effect on the rate of recovery from inactivation. Mean results ± SEM of 13 cells from 4 animals (left). Representative recordings of I_Na_ at +12°C and +25°C.

## DISCUSSION

The dissociation of atrial and ventricular beating rates at the *T*_BP_ of *f*_HV_ corroborated our hypothesis (ii) about the temperature dependent failure of ventricular excitation. The sharp increase of *f*_HA_ above the *T*_BP_ of *f*_HV_ suggests that the beating rate of the primary pacemaker and the ability of the atrium to follow it are not impaired at temperatures above the upper critical temperature of the rainbow trout (+25.6-26.9°C) (Hokanson et al., 1977; Kaya, 1978; Ekström et al., 2014; Gilbert et al., 2019). In contrast, the ventricle can’t keep pace with the pacemaker rate and therefore the heart has two separate beating rates at critically high temperatures, one for the atrium and another for the ventricle. The nearly exponential increase and the high peak value of *f*_HA_ in rainbow trout (188 pbm at +27°C) matches well with the thermal response of AP rate in the isolated pacemaker cells of the brown trout (193 bpm at +26°C) (Haverinen et al., 2017). Collectively, these findings indicate that the temperature-induced bradycardia is specific for the ventricle, and strongly suggest that the high temperature induced depression of cardiac output in fish (Gollock et al., 2006; Steinhausen et al., 2008; Clark et al., 2008; Mendonca and Gamperl, 2010; Ekström et al., 2016) is due to the inability of the ventricle to follow the sinoatrial rate. Our results are consistent with the early *in vitro* findings of both teleost and elasmobranch hearts, in that the ventricle is the most heat sensitive cardiac compartment of the fish heart, which loses the ability to follow the increasing beating rate of the upper cardiac compartments (Koehnlein, 1933; Markowsky, 1933).

In the normally functioning vertebrate heart, P wave always precedes QRS complex. In the +12°C-acclimated rainbow trout, atrial P wave and ventricular QRS complex became partially dissociated at about +25°C. This is a functional AV block and the cause for the temperature-induced ventricular bradycardia. In all 4 trout where P was clearly visible, at some point of warming every other ventricular beat was missed causing a 2:1 AV block. With further warming this progressed to a 3:1 AV block, which explains the deepening ventricular bradycardia above *T*_BP_ of *f*_HV_. In the human heart, AV block is classified in three categories (Vogler et al., 2012). In the first-degree AV block, each QRS complex is preceded by P wave. The only deviation from the normal heart function in this benign condition is the slowed AP conduction (prolongation of PR interval) in the AV node. Characteristic for the second-degree AV block is the intermittent block of AP propagation between the atria and the ventricles i.e. occasional dissociation of P wave and QRS complex. Based on ECG morphology, two types of the second-degree AV block, Mobitz type 1 and Mobitz type 2, are separated. The type 1 block is caused by the malfunction of the AV node which results in non-conducted P waves. ECG of the type 1 block is characterized by a prolonged PR interval i.e. there is a gradual slowing of AP conduction with successive beats in the AV node which then results in a skipped ventricular beat. The type 2 block also appears as dissociation of P wave and QRS complex but without the conduction delay in the AV node. In the type 2 block, the failure occurs in the ventricular conducting system and is often associated with an increase in QRS duration. The AV block of the trout heart resembles the human second-degree type 2 block in that P wave and QRS complex are dissociated but without prolongation of the AV conduction time. In fact, PQ interval was gradually shortened and there was slight increase in QRS duration when the fish was acutely warmed. Similarly, in tench (*Tinca vulgaris*), eel (*Anguilla vulgaris*) and torpedo (*Torpedo ocellatta* and *T. marmorata*) conduction time between atrium and ventricle shortened at temperatures between +5°C and +25°C (for tench and eel) and +2°C and +28°C (for torpedo). However, at temperatures between +25°C and +30°C conduction started to slow down, and this was considered to cause AV block and dissociation of atrial and ventricular contractions (Markowsky, 1933). Our present findings *in vivo* (ECG) and those of Markowsky and Kohnlein *in vitro* (contractile studies) are closely similar, but the explanations of the phenomenon differ. While Markowsky and Kohnlein favour the failure of impulse conduction in the AV canal as a causal factor, we are suggesting that it is the increased excitation threshold of ventricular myocytes which precludes the ventricular contraction. In fact, both factors could be involved. It has been noticed in the sinoatrial pacemaker cells of the brown trout that high temperature does not depress AP frequency but it reduces shortens the duration and reduces the amplitude of the nodal AP (Hassinen et al., 2017). If the same holds for the nodal cells of the AV canal, the ventricular failure could be partly related to the small APs of the AV nodal cells and partly caused by increased excitation threshold of ventricular myocytes. APs are conducted through the AV canal, but they are too small and short to trigger APs in ventricular cells whose V_th_ is elevated (Note: the strength-duration relationship shows that short pulses require a larger current to trigger AP). However, little is known about the temperature induced changes in excitability of the fish AV canal. Clearly, the role of the AV canal in thermal responses of the fish heart needs to be examined.

It has been suggested that the warming induced bradycardia in fish could be an adaptive reaction to high temperature by improving myocardial oxygenation (Farrell, 2007; Ekström et al., 2014; Ekström et al., 2019; Gilbert et al., 2019). Mechanistically this could happen via the release of acetylcholine from the vagal nerve endings and subsequent hyperpolarization of the maximum diastolic potential of the pacemaker cells, if the cholinergic tone was increased by acute warming (Campbell et al., 1989). However, the findings on the cholinergic regulation of the thermal tolerance of *f*_H_ in rainbow trout are contradictory. Some studies suggest that blocking of the cholinergic nervous transmission has no effect on the temperature tolerance of *f*_H_ (Ekström et al., 2014; Ekström et al., 2019), while others suggest that temperature tolerance is decreased by the cholinergic block (Gilbert et al., 2019). Both conclusions are based on the beating rate of the ventricle. The present findings show that *f*_HA_ and *f*_HV_ become separated when *f*_H_ is about 120 bpm. In the light of the present study, the 2-degree depression in *T*_BP_ of *f*_H_ in the presence cholinergic block (Gilbert et al., 2019) is explained by the slightly higher *f*_H_ of the atropine treated fish i.e. the “cut-off” frequency of the AV block (about 120 bpm) is achieved at slightly lower temperature. Thus, the cholinergic tone *in vivo* can indirectly increase *T*_BP_ of *f*_HV_ by shifting the cut-off frequency of the AV block to slightly higher temperatures. However, the strong increase of *f*_HA_ above the *T*_BP_ of *f*_HV_ indicates that the sinoatrial rate remains intact and therefore excludes the cholinergic “brake” as an explanation for the temperature-induced bradycardia. To conclude, the present findings show that the temperature induced ventricular bradycardia is a non-adaptive trait, because it is caused by malfunction of the cardiac excitation system.

At the cellular level, the reduced ventricular excitability was evident as temperature-induced increase in CD and I_th_. With acute warming both V_rest_ and V_th_ become more negative but the effect of temperature on V_rest_ was stronger than on V_th_, and therefore the voltage distance between the two (CD) increased: more current (I_th_) was needed to trigger AP. According to the TDEE hypothesis, the failure of ventricular excitation is due to the discordant temperature-dependencies of I_Na_ and I_K1_, the two opposing currents decisive for the initiation of ventricular AP (Varghese, 2016; Vornanen, 2016). Consistent with the TDEE hypothesis the depression of EE was associated with an increase in the outward I_K1_ of the resting ventricular myocyte and a decrease in the charge transfer by I_Na_ of the active myocytes. Collectively these findings indicate that a mismatch between the inward I_Na_ and the outward I_K1_ develops at high temperatures, which results in depressed excitability of ventricular myocytes (Vornanen, 2016). This was further tested using the classical method to measure EE of cells and tissues i.e. to determine the strength-duration relationship (Weiss, 1901; Lapicque, 1909; Geddes, 2004). This analysis gives two variables (1) rheobase which is the minimum stimulus strength of an indefinite long depolarizing pulse needed to trigger AP, and (2) chronaxie which is the minimum duration of stimulus pulse with a twice the rheobase strength sufficient for initiation of AP (Irnich, 1980). In the space clamped ventricular myocytes, the rheobase current was 56.6% larger at +25°C than at +12°C indicating an increased demand for depolarizing current/charge. This is due to the increased outward leakage of K^+^ ions as indicated by the lower Rin and the higher I_K1_ at +25°C relative to +12°C. K^+^ efflux of the quiescent myocyte shunts the inward current inflow from the upstream cell with the consequence that the diminished inward charge transfer by I_Na_ cannot depolarize the membrane to the V_th_. Basically the present results are consistent with the earlier findings on another teleost species, the brown trout (*Salmo trutta fario*) (Vornanen et al., 2014).

In multicellular cardiac tissue, the increased outward K^+^ leakage of the quiescent myocytes forms the current sink of the downstream myocyte, which must be exceeded by the inward I_Na_ from the activated upstream myocyte or the current source (Varghese, 2016). Robust excitability of the ventricle requires that there is some excess of the inward I_Na_ relative to the outward I_K1_. Owing to the surplus of I_Na_ small changes in physiological or environmental conditions are unlikely to cause failure of ventricular excitation. The excess of the inward current relative to the outward current is called safety factor or safety margin, which can be defined as the ratio between charge provided by the upstream source cell and charge demanded by the downstream sink cell for excitation (Shaw and Rudy, 1997). When the safety factor is ≥1.0 excitation is ensured and if the safety factor is <1.0 excitation will fail. At high temperatures the increase of I_K1_ with the simultaneous decrease of I_Na_ charge transfer “consumes” the surplus of the source current resulting in source-sink mismatch and failure of the ventricular excitation. Notably, in rainbow trout ventricular myocytes the charge transfer via I_Na_ started immediately to decline when temperature exceeded +12°C. At the breakpoint temperature of the *f*_HV_ the charge transfer by I_Na_ was reduced by about 46% and the membrane leakage was approximately doubled. From these changes it can be estimated that the safety factor at the acclimation temperature of the fish (+12°C) was more than 2.0. At +25°C, the safety margin is probably consumed (<1.0) resulting in bradycardia. The main reason for the decline of the charge transfer by I_Na_ is the temperature-induced increase in the rate of current inactivation. Also, the rate of current activation is slightly increased by high temperatures but its effect on charge transfer is less than that of inactivation, because the activation of I_Na_ is almost instantaneous already at +12°C and therefore not much affected in absolute terms by warming.

## CONCLUSIONS

AV block due to the mismatch between I_Na_ source current and I_K1_ sink current (membrane leak) of ventricular myocytes provides a mechanistic explanation for the high temperature-induced ventricular bradycardia. While the source-sink mismatch concept is consistent with the oxygen and capacity limited thermal tolerance – a hypothesis that oxygen delivery by circulation and ventilation set the thermal tolerance limits of ectotherms (Pörtner, 2001) - its scope of application might not be limited to cardiac function. The basic principles of EE and the sourcesink mismatch hypothesis as presented here for the rainbow trout ventricular myocytes should be valid for other electrically excitable tissues and cells (nerves and muscles) as well (Koester and Siegelbaum, 2000). Therefore, it may be premature to assume that the temperature-dependent weakening of physiological performance of the fish is solely due to the impairment of cardio-ventilatory function. It is equally possible that simultaneously (or at lower temperatures) with the cardiac dysfunction sensory and motor functions and coordination of the body homeostasis of the fish are also compromised due to the source-sink mismatch in neuronal networks and muscle tissues.

## List of abbreviations

AP: Action potential
APD_50_: Action potential duration at 50% of repolarization
CD: Critical depolarization
C_m_: Membrane capacitance
ECG: Electrocardiogram
EE: Electrical excitability
*f*_H_: Heart rate
*f*_HA_: Beating rate of atrium
*f*_HV_: Beating rate of ventricle
I_th_: Threshold current
R_in_: Input resistance
*T*_BP_: Breakpoint temperature
V_th_: Threshold potential
V_rest_: Resting membrane potential

## Acknowledgements

We would like to thank Kontiolahti fish farm for providing the fish. Anita Kervinen is acknowledged for taking care of the fish at the university and preparing solutions for experiments.

## Author contributions

conceptualization: M.Vornanen

Investigation: J.Haverinen

Original draft: M.Vornanen and J.Haverinen

Writing - review & editing: M.Vornanen

Visualization: J.Haverinen

Funding acquisition: M.Vornanen

